# Development of self-healing hydrogels to support choroidal endothelial cell transplantation for the treatment of early age related macular degeneration

**DOI:** 10.1101/2024.06.07.597936

**Authors:** Narendra Pandala, Ian Han, Kevin Tobin, Nicole Brogden, Kelly Mulfaul, Robert Mullins, Budd Tucker

**Author notes:** **Corresponding author:** Budd A. Tucker.

## Abstract

In retinal diseases such as age-related macular degeneration (AMD) and choroideremia, a key pathophysiologic step is loss of endothelial cells of the choriocapillaris, the dense vascular bed required for maintaining health and function of the retina. As such, repopulation of choroidal vasculature early in the disease process may halt disease progression. Prior studies have shown that injection of donor cells in suspension results in significant cellular efflux and poor cell survival. As such, the goal of this study was to develop a hydrogel system designed to support CEC transplantation. A library of hydrogels was synthesized using laminin (i.e., LN111, LN121, and LN421), carboxy methyl chitosan, and oxidized dextran via reversible Schiff base chemistry. Each of the developed hydrogels was readily injectable into the suprachoroidal space, with excellent gelation, mechanical, and degradation properties. Laminin-based hydrogels were compatible with immortalized CEC survival in vitro, and suprachoroidal injection of LN111 and LN121 containing gels were well-tolerated in an in vivo rat model, whereas LN421 containing gels caused significant chorioretinal inflammation. Hydrogels were detected in the suprachoroidal space of immunosuppressed rats at 1-week post-injection and were completely resorbed by 1-month post-injection. There were significantly more CECs retained in immunosuppressed rats that received cell laden hydrogels than those that received unsupported cell suspensions (i.e., CECs only). These findings pave the way for future CEC replacement studies in animal models of choroidal cell loss toward the development of future therapies.

**Statement of significance:** Age related macular degeneration (AMD) is a leading cause of untreatable blindness in the industrial world. A key pathologic step in AMD is loss of the choriocapillaris endothelial cells, which provide vascular support to the overlying retina, including the light-sensing photoreceptors. We believe that choroidal cell replacement early in disease may prevent retinal cell death and subsequent vision loss. In this study, we present a strategy for repopulating the choriocapillaris using choroidal endothelial cell laden hydrogels. Specifically, we demonstrate the synthesis and characterization of 3 different laminin-based hydrogel systems. LN111 and LN121 hydrogels were found to increase retention of choroidal endothelial cells following suprachoroidal transplantation. These findings pave the way for future cell replacement studies in animal models of choroidal cell dropout.

## Introduction

Age related macular degeneration (AMD) is a genetically complex progressive disorder characterized by death of choroidal endothelial, retinal pigment epithelium (RPE), and photoreceptor cells, resulting in irreversible vision loss. AMD can be subdivided into two broad categories based on the presence or absence of aberrant blood vessel growth: 1) wet or neovascular AMD, characterized by choroidal neovascularization, and 2) dry or non-neovascular AMD, with geographic atrophy representing end-stage disease. While numerous effective treatments designed to prevent growth of new blood vessels exist for patients with wet AMD, limited treatments exist for patients with dry AMD. In earlier stages of dry AMD, micronutrient supplementation (known as the AREDS formulation) [1, 2] may decrease the rate of progression from dry to wet AMD [3, 4]); however, such treatment does not slow choroidal or photoreceptor cell loss. Recently, a C3 inhibitor (pegcetacoplan)[5, 6] and a C5 inhibitor (avacincaptad pegol)[7, 8] were approved by the FDA for the treatment of geographic atrophy secondary to AMD[9]. Because these treatments are currently indicated only for geographic atrophy (i.e., after significant complement induced injury and vascular loss have already occurred), the impact on slowing the progression of disease is modest.[10]

Choriocapillaris endothelial cell (CEC) death, which typically precedes loss of RPE and photoreceptor cells [11–13], results in the formation of non-functional “ghost” vessels[14]. As the choriocapillaris is the primary means by which waste material is removed from the overlying retina, and ghost vessels are most frequently observed directly beneath drusen deposits[15] (i.e., a buildup of cellular biproducts between the RPE and Bruchs membrane), it is compelling to believe that choroidal cell death is an inciting factor in drusen deposition, RPE cell dysfunction, and photoreceptor cell loss. This is further supported by the fact that leading genetic risk factors for development of AMD are associated with the complement cascade and the choriocapillaris is the primary site of complement deposition [16]. For these reasons we hypothesize that repopulation of the choriocapillaris with functional choroidal endothelial cells early in disease may prevent drusen deposition, RPE cell dysfunction, photoreceptor cell death, and subsequent loss of vision.

To repopulate the choriocapillaris, one could envision injecting cells directly into the capillary bed beneath Bruch’s membrane (i.e., the supportive elastic layer under the RPE). Unfortunately, unlike the neural retina, which can be readily detached from the RPE creating a space into which cells can be delivered, no such virtual space exists between the RPE and choroid, and a transretinal approach of injection would require a retinotomy. As injection of therapeutic cells into the circulatory system is undesirable, the most logical option for choroidal cell replacement is to inject donor cells into the suprachoroidal space. The suprachoroidal space is readily accessible via injection to separate the sclera from the choroidal vasculature and is a viable location for choroidal drug delivery [17]. Microneedles designed to penetrate the sclera and deliver product directly to the choroid exist and are being used clinically [18]. Unfortunately, injection of cells through a fine gauge microneedle in the absence of structural support, has been associated with significant shear stress induced cell death [19], cellular efflux post-needle removal [20], and poor donor cell retention[21–23]. To enable choroidal cell retention following suprachoroidal injection novel materials that are designed to address these issues will be required.

Hydrogel based biomaterials, which have been widely used in tissue engineering applications to enhance delivery of drug, protein, and cellular products [21–23], have the potential to enhance survival and retention of choroidal endothelial cells following suprachoroidal injection. Unlike solid cell support scaffolds, self-healing hydrogels can be injected through small gauge cannulas and reform within the injection site. While they reduce sheer stress and hold donor cells within the injection site immediately following delivery, biomaterial-based hydrogels are typically short lived and allow cells to migrate away from the injection site into their desired location upon resorption.

In this study, we report development of a library of injectable biomaterial-based hydrogels using laminin, chitosan, and dextran via amine aldehyde reversible Schiff base chemistry. Laminins were chosen as one of the primary components of these hydrogels as they are extracellular matrix (ECM) proteins that naturally reside in the CNS where they promote cellular adhesion and survival via integrin activation [24–29]. Likewise, chitosan and dextran have been used in many regenerative medicine applications with excellent biocompatibility [30–36]. Self-healing hydrogel formulations with one of three different isoforms of human laminin at varying concentrations were synthesized and compared. We evaluate the tolerability of these hydrogels in vitro, as well as via suprachoroidal delivery in an immunosuppressed rat model in vitro. We compare the post-transplant CEC donor cell survival and retention across the formulations tested. These findings pave the way for future CEC replacement studies toward the development of AMD treatment.

## RESULTS

### Hydrogel synthesis and characterization

Self-healing hydrogels were synthesized by utilizing the Schiff base reversible reaction between aldehydes obtained from oxidized dextran and free amines obtained from chitosan and laminin (**Figure 1A**).

**Figure 1:**
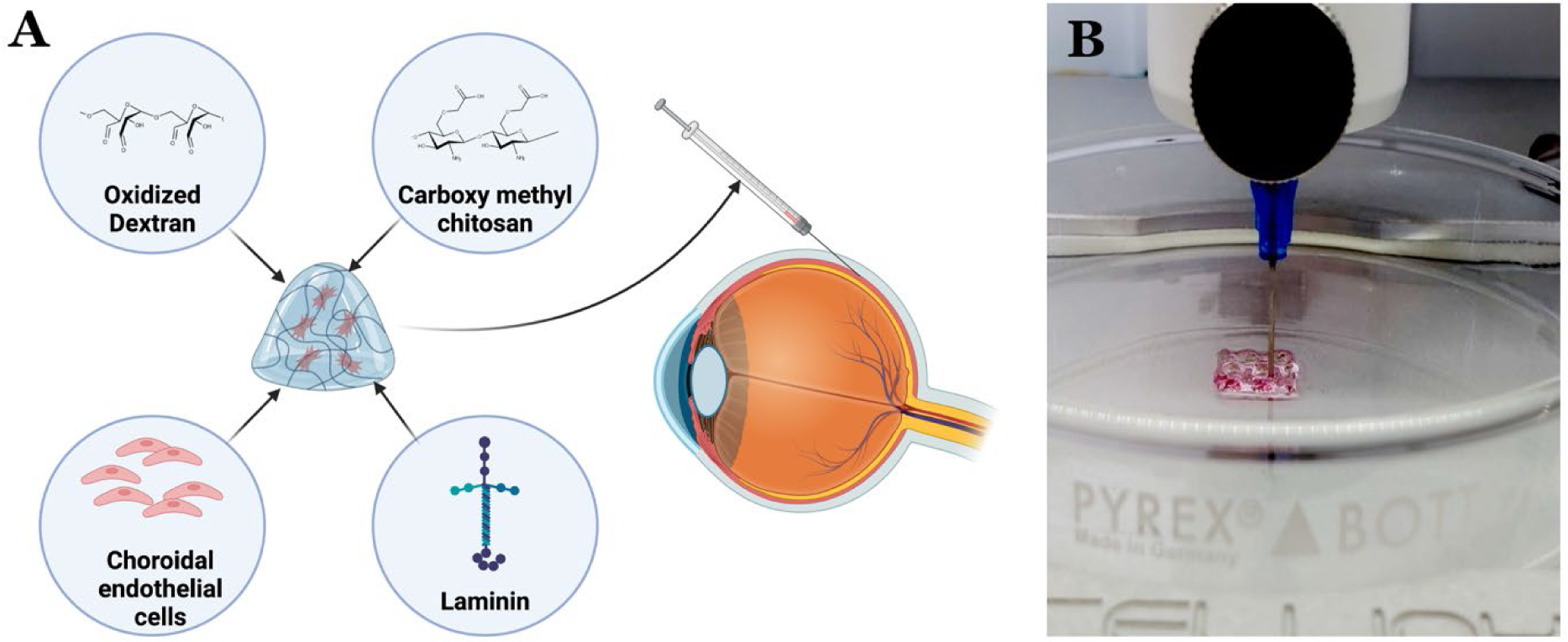
A) Schematic showing the gel fabrication and suprachoroidal injections. B) Gel-30-LN 111 being extruded through a 23G needle using a 3D bioprinter (BioX, Cellink). The hydrogel was set at 37°C for 2 hours before extrusion.

Dextran was oxidized using sodium periodate, which was confirmed via ^1^H NMR (**Figure S1**). To maintain the hydrogels at physiological pH and increase chitosan solubility, we used carboxymethyl chitosan for hydrogel fabrication [37, 38]. By using carboxymethyl chitosan, oxidized dextran, and three different isoforms of laminin (namely LN111, LN121, and LN421), we synthesized a library of seven independent hydrogels (**Supplementary Table 1**). Hydrogels were synthesized by altering the concentration of laminin and chitosan while keeping the amount of dextran constant. The gels are labelled Gel-0, Gel-30, and Gel-50, representing increasing concentrations of Laminin (0, 30, and 50 µg/ml respectively). We attempted to generate hydrogels in the absence of chitosan, that contained laminin (80 µg/ml) and oxidized dextran only. As these gels were not consistent (i.e., partial gels with liquid) they were excluded from our analysis. To demonstrate the injectability of the synthesized hydrogels, we loaded the hydrogel precursors into a syringe and extruded them using a bioprinter (**Figure 1B**). Hydrogels were easily extruded at a pressure of 50kPa and rapidly formed a cohesive mass, demonstrating both their injectability and self-healing properties.

### Mechanical characterization

Small amplitude oscillatory shear (SAOS) rheometry was used to determine the gelation time of each hydrogel. Gel-0-LN111 had a gelation time (G’>G”) of approximately 16 mins, Gel-30-LN111 had a gelation time of approximately 21 mins, and Gel-50-LN111 had a gelation time of approximately 29 mins (**Figure 2A-C, Figure S2**). While gelation time increased as the concentration of laminin increased, gelation time was independent of laminin isoform (i.e., there was no significant difference in gelation time between LN111, LN121, and LN421 containing gels at the same laminin concentration). Storage moduli of each hydrogel was determined using frequency sweeps (**Figure 2 D-F, Figure-S3**). Interestingly, a similar inverse trend was observed. Specifically, as the amount of laminin increased, the storage modulus of the hydrogel decreased. Again, this effect was independent of the laminin isoform used (i.e., there was no significant difference in storage modulus between gels containing the same concentration of LN111, LN121, or LN421).

**Figure 2:**
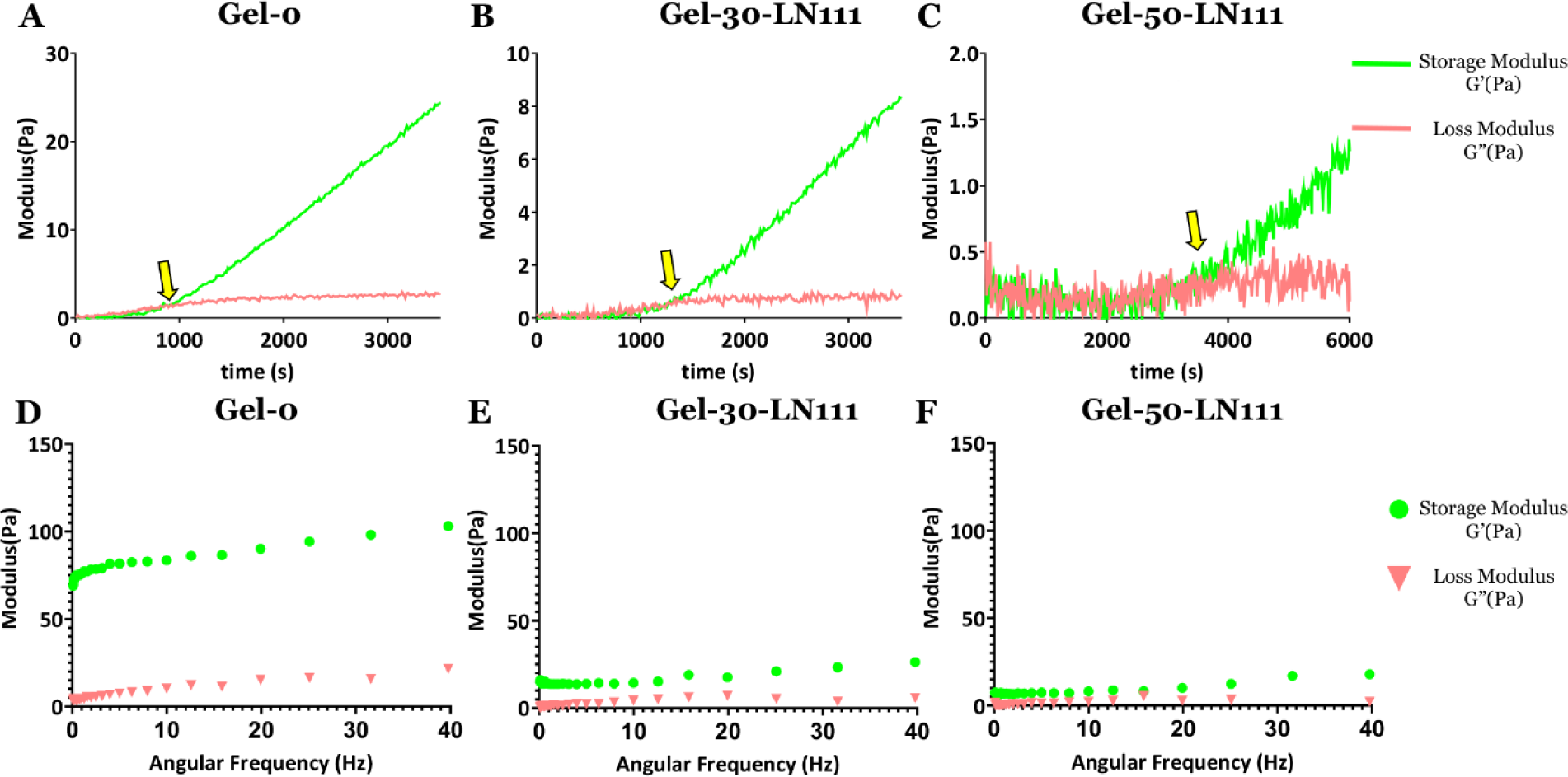
Time sweeps of (A) Gel-0 (0 µg/ul-laminin), (B) Gel-30 (30 µg/ul-LN111) and (C) Gel-50 (50 µg/ul-LN 111), showing the gel formation time at the crossover point (G’>G”). Frequency sweeps of the precast (D) Gel-0, (E) Gel-30, and (F) Gel-50, fabricated using LN111, showing the frequency dependence of the storage modulus (G’) and loss modulus (G”). Pa = Pascals.

### In vitro biocompatibility of the gel system

To evaluate biocompatibility of each of the developed hydrogels in vitro, our previously reported human choroidal endothelial cell line (ICEC2-TS) was used [39, 40]. This line was generated from primary human endothelial cells and expresses markers such as CDH5, CD34, vWF, and PECAM1. As these cells (which were immortalized via SV40 T antigen) are able to form vascular tubes when cultured in Matrigel and reintegrate into decellularized human choroids in the dish [40], they are ideal for developing the first pass for novel hydrogel mediated cell delivery approaches. To determine if the developed hydrogels were compatible with ICEC2-TS cells, fluorescent live dead assays were performed. Specifically, gel precursors and ICEC2-TS cells were mixed, and cellular survival was analyzed using calcein-AM (green - live cell stain) and ethidium homodimer-1 (red - dead cell stain). Gels were placed in cell culture media at 37C and imaged at 1- and 7-days later (**Figure 3 & Figure S4**). A very low number of dead cells was detected in either condition, indicating that each of the developed gels had excellent biocompatibility in vitro. Cells were spherical and remained in suspension for up to 7-days. Beyond 7-days, gels began to degrade, upon which cells were liberated, adhered to the culture plates, and began to spread across the cell culture surface (**Figure 3**). These degradation profiles are similar to those reported for similar gels [41, 42].

**Figure 3:**
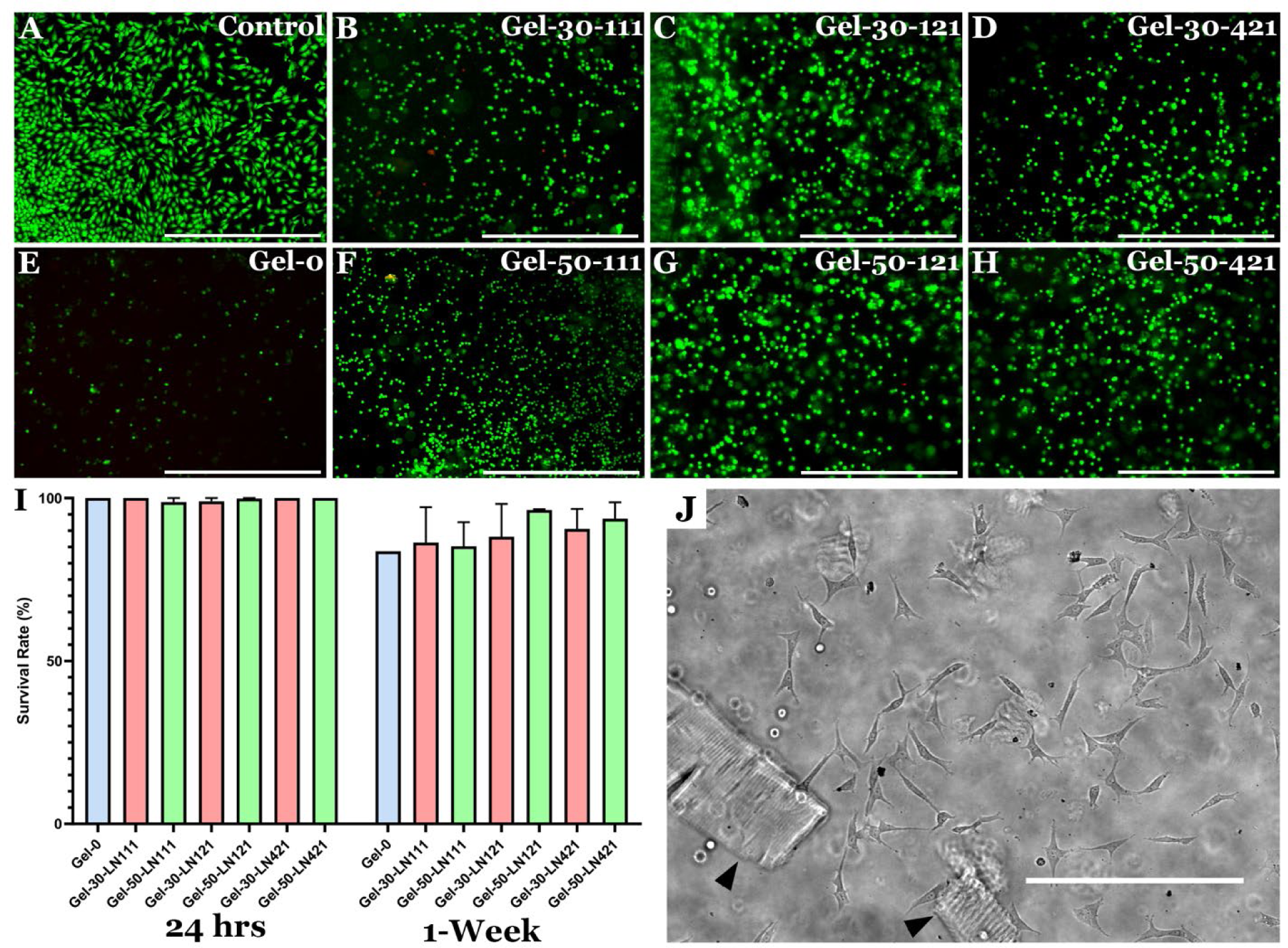
Live dead staining of cell laden hydrogels synthesized using 3 different isoforms of Laminin (day-1). Human primary CECs were stained using a live dead stain consisting of calcein-AM as the live stain (green) and ethidium homodimer-1 as the dead stain (red) (Scale bar is 1000µm)

To evaluate the effect of injection induced shear stress on cell viability, cell laden gels were injected through a 30G canula and cell survival was evaluated (**Figure S5**). Regardless of laminin isoform used, Gel-50 passed through the canula with the least amount of resistance and greatest cell viability. For this reason, only Gels with 50µg/ml of laminin were used in subsequent in vivo studies.

### In vivo hydrogel assessment

To evaluate biocompatibility of each of the developed hydrogels in vivo, we injected them into the suprachoroidal space of wild type Sprague Dawley rats. At one-week post-injections the eyes were examined via indirect ophthalmoscopy, fundus photography (**Figure 4A-D**) and optical coherence tomography (OCT) (**Figure 4E-H**). The retina looked intact without overt signs of damage detectable via fundus photography. Mild retinal hyperreflectivity (**Figure 4E yellow arrow**) and thickening of photoreceptor cell outer segments (**Figure 4G yellow arrow**) with otherwise intact retinas was observed via OCT. In vivo biocompatibility was further analyzed via immunofluorescence staining of paraformaldehyde fixed and cryo-sectioned eyes using antibodies targeted against the immune cell marker IB4 (**Figure 4I-L**). Interestingly, LN421 hydrogel injected eyes had an elevated inflammatory response, drawing an increased number of macrophages to the injection site than either of the other two hydrogels (LN 111, 121) or vehicle injection control (**Figure-4I-L**). This might be because of the presence of LN111 and LN121 natively in the ECM of RPE cell basement membranes [43, 44], whereas LN 421 is usually produced by the mesangial cells of the kidney [45, 46]. Going forward, only LN111 and LN121 containing hydrogels were used for subsequent cell injection analysis.

**Figure 4:**
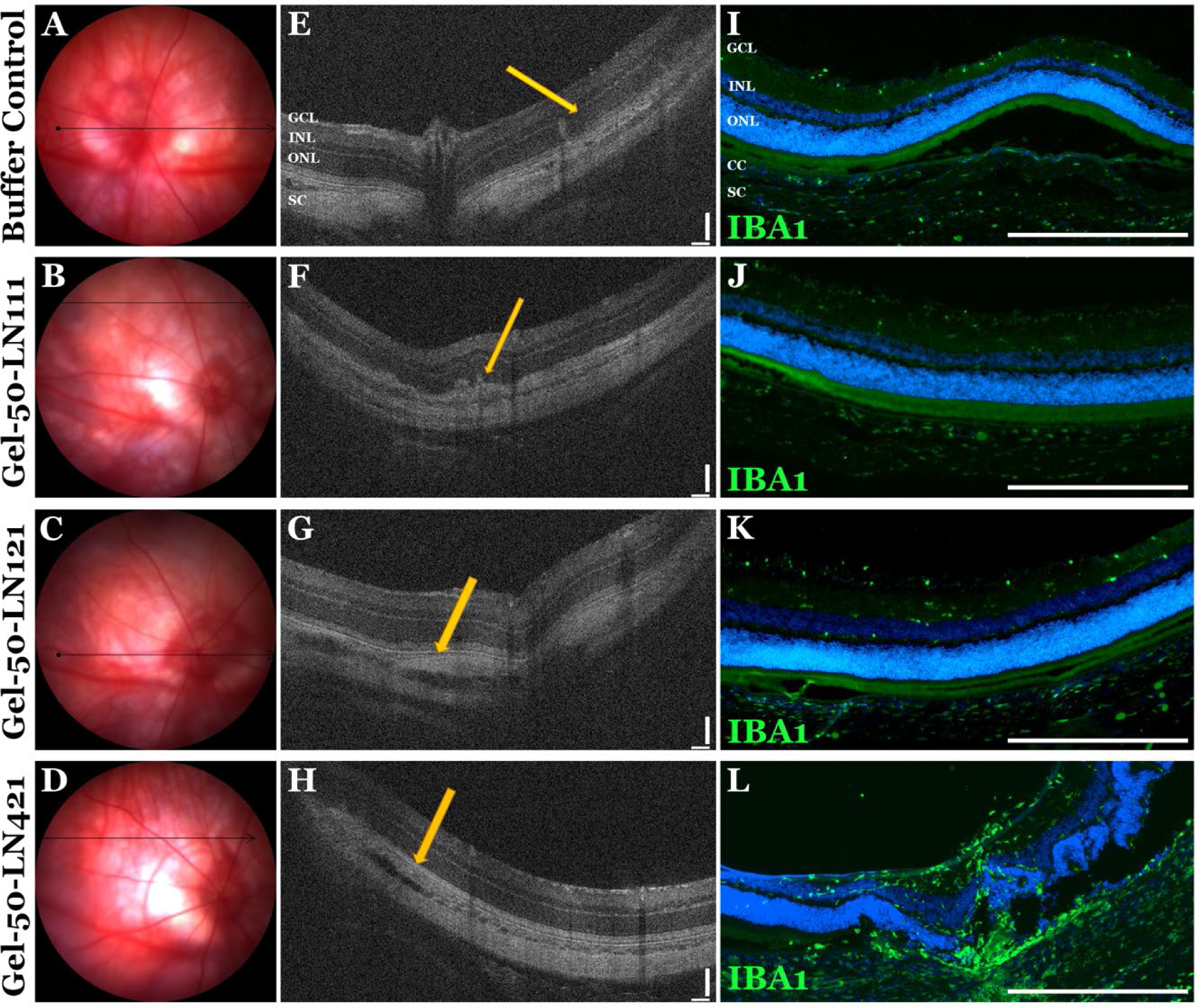
**A-H)** Fundus photographs (A-D) and OCT images of SD animals at 1 week following suprachoroidal hydrogel injection. Complete media was used as a vehicle control. Yellow arrow in panel E is showing a region of mild retinal hyperreflectivity. Yellow arrow in panel G is showing a region of mild thickening of the photoreceptor outer segments. Yellow arrow in panels F and H are pointing to the injection site, which is observed by the suprachoroidal cleft. **I-L)** Fluorescent images showing macrophage/microglial cell infiltration (as determined by IBA1 staining) in eyes that received a suprachoroidal dose of either hydrogel or vehicle control. (Scale bar for I-L is 100µm).

To determine if the cell containing hydrogels were well tolerated, LN111 and LN121 hydrogels laden with choroidal endothelial cells were injected into the suprachoroidal space of immunocompromised SRG (Sprague-Dawley Rag2/Il2rg double-knockout) rats (Charles River, Wilmington, MA, USA). To evaluate ocular histology following suprachoroidal injection, eyes were embedded in Paraffin, sectioned, and subjected to H&E staining. At 4-weeks post-injection hydrogels were completely resorbed. The H&E sections show an intact retinal layer without any damage (**Figure-5A-C**).

## DISCUSSION

Hydrogels produced using amine aldehyde chemistry have been widely characterized and demonstrated to have favorable biocompatibility profiles [47–50]. By utilizing amines in hydrogel precursors, it is possible to augment small molecules, peptides, or proteins via Schiff base chemistry [51]. For instance, extracellular matrix molecules known to promote cellular attachment and survival can be readily incorporated. As indicated above, the purpose of this study was to develop a biomaterial-based hydrogel system to enhance choroidal endothelial cell retention following suprachoroidal transplantation. To achieve this goal, we generated a library of novel self-healing hydrogels that contained chitosan, dextran, and one of three different isoforms of laminin.

Chitosan is a linear polysaccharide, that has been reported to have anti-inflammatory[52], antimicrobial[53], antioxidant[54] and wound healing[55] properties, making it an excellent candidate for a variety of tissue engineering applications [31]. However, as chitosan is soluble at low pH, non-crosslinked precursors solutions are not typically biocompatible (i.e., acidic precursor solutions are cytotoxic). To overcome this hurdle, carboxy methylated chitosan, which can be maintained in solution at physiologic pH, was used [37, 38]. Carboxy methylated chitosan prepolymers can be formulated in growth media, PBS, or other cell compatible buffers to facilitate the Schiff base reaction without negatively impacting cell viability [56]. Dextran, like chitosan, is an inert polysaccharide that can be readily functionalized for hydrogel production [30, 32]. Dextran’s modified with either methacrylate, vinyl sulfones, or aldehydes, have been used in a variety of hydrogel formulations [33–36]. We oxidized the hydroxyl groups on the dextran to aldehydes, using the sodium periodate-based oxidation technique because of the simplistic reaction procedure and purification steps. We chose this modification to facilitate the Schiff base chemistry, which in turn is highly biocompatible and does not produce any cytotoxic byproducts [47–50].

Unlike chitosan and dextran, laminins are extracellular matrix glycoproteins that consist of α, β, and γ polypeptide chains. Laminins, which bind and activate integrin receptors in the plasma membrane, are used routinely to promote cellular adhesion, migration, and survival both in vitro and in vivo [24, 25, 57-61]. Here, we describe the production of hydrogels containing either LN111, LN121, or LN421. LN111 is a major component of Matrigel, a mixture of extracellular matrix proteins isolated from mouse EHS tumor cells, which is widely used in cell culture to promote cellular attachment and differentiation[62]. At 37°C Matrigel forms a solid 3-dimensional gel into which seeded endothelial cells can form dense vascular networks[63, 64]. In addition, LN111 has also been identified as a component of retinal pigment epithelial cell basal lamina and Bruch’s membrane[65], making it an obvious choice for choroidal endothelial cell delivery.

While ocular expression of full length LN121 has not been reported, each of its individual polypeptide chains (i.e., α1, β2, and γ1) have been identified. For instance, β2 laminin was shown to be expressed during early retina genesis[44]. Loss of beta 2 laminin induced asymmetric cell division, causing early depletion of the retinal progenitor cell pool, overproduction of rod photoreceptors and a reduction in the number of late born bipolar and muller glial cells[44]. Interestingly, compared to LN111, LN121 was found to have a higher affinity for integrin binding [66] and more effective at promoting neurite outgrowth in vitro[66].

Unlike LN111 and LN121, which were selected based on their importance in both the developing and adult retina, LN421, which is predominantly expressed in the kidney, was selected because of its reported role in endothelial cell fate commitment[67], survival [68], and microvascular repair [45, 46]. Interestingly, of the 3 laminin isoforms tested, hydrogels containing LN421 were the least well tolerated in the suprachoroidal space. Specifically, injection of LN421 containing hydrogel resulted in increased microglial cell activation and/or macrophage infiltration as determined by IBA1 staining. Interestingly, this appeared to be independent of a direct toxic effect, as cells cultured in LN421 containing hydrogels did not differ in morphology or viability from those cultured in LN111 or LN121 containing hydrogels (i.e., Figure 3). Of note, unlike previously reported hydrogels containing RGD peptides, which in some cases promoted cellular spreading within the gel [33–36], choroidal endothelial cells cultured in the reported laminin based hydrogels remained immobilized yet viable within the hydrogel network until they were released via gel degradation. As our intent was to hold cells in place for several days to ensure wound closure and optimal donor cell retention, this lack of cell adhesion was advantageous for our intended application.

When designing an injectable hydrogel for cell delivery, the storage modulus of the material must be considered. Hydrogels with lower storage moduli are typically easier to inject [69]. This is particularly important when injections are performed using small gauge cannulas, which is often the case with ophthalmic microsurgery. As the inner diameter of the injection cannula decreases, the pressure and in turn shear stress experienced by the cells contained within the cannula increases. This is important as sheer stress encountered during the injection process is thought to be a major contributor to poor cell viability following transplantation [19]. Given the low concentrations of our hydrogel precursor solutions, all of the gels developed in this study were found to have a low storage modulus as determined via rheometry. Of the tested hydrogels, Gel-50 was found to have the lowest storage modulus regardless of laminin isoform used. While we did not detect a significant increase in cell viability using this formulation, this was not entirely surprising given that cell viability following injection was >95% with the highest storage modulus (i.e., reducing storage modulus further would have little room for improvement).

In addition to storage modulus, gelation time must also be considered when designing a hydrogel for cell delivery. If the hydrogel solidifies too quickly it can be challenging to load and expel from the injection device. This is especially important for cell replacement studies in which the injection devices are loaded in a biological safety cabinet in the lab before being transported to the surgical suite for ocular injection. Hydrogels with rapid gelation times often solidify within the injection device, clogging the injection canula and making it difficult to achieve consistent results. For hydrogels with very slow gelation times, viability of cells contained within the prepolymer mixture can be negatively impacted. Our goal in this study was to develop a prepolymer formulation that would gel within 60-90 minutes of being mixed with the donor cells. This would give us ample time to prepare the surgical suite, anesthetize the animals, load the injection devices and transport them to the surgical suite. Our goal was to have the material gel sufficiently within the injection device to hold the cells in place once injected but still soft enough to allow the gel to be injected without clogging the canula. Interestingly, we found that increasing the concentration of laminin contained within each hydrogel lowered both the gelation kinetics and moduli, indicating that it induced favorable structural changes within the hydrogel network.

One additional consideration when designing hydrogels for suprachoroidal endothelial cell injection is degradation time. As demonstrated here, once solidified, the developed hydrogels did an excellent job of immobilizing choroidal endothelial cells without having a negative impact on viability. It was not until the polymers began to degrade that the cells were liberated (Figure 3). Our goal was to produce a gel that would remain intact, holding cells long enough to allow the surgical wound to heal (preventing cellular efflux), then degrade, liberating cells for integration with the native tissue. We reasoned that having the hydrogel completely resorbed within 1 week following injection would be ideal. With daily media changes in vitro, significant degradation was detected by 7-days. Interestingly, as the concentration of laminin increased the degradation time decreased suggesting that higher concentrations of laminin were preferable. As hydrogels were not detectable in any animal at 4-weeks post-injection (Figure 5), these findings suggest that the in vitro and in vivo data were highly corelated and consistent with similar published dextran and chitosan-based hydrogel systems [41, 42].

**Figure-5:**
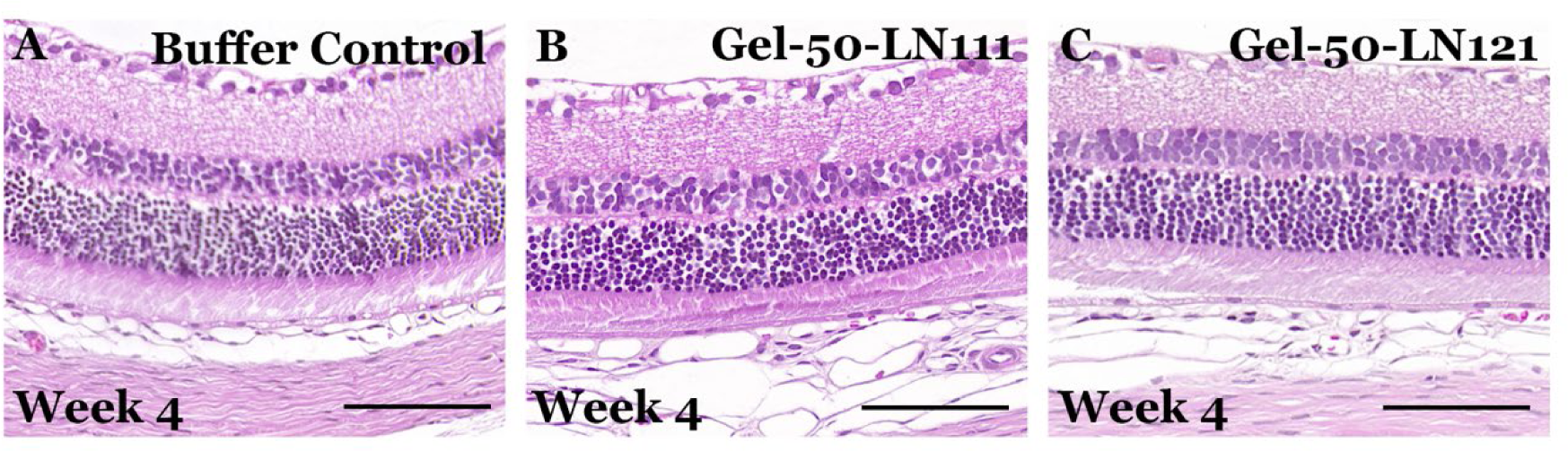
**A-C)** H&E stain demonstrating complete hydrogel resorption and intact retina at 4-weeks following suprachoroidal injection. (A) Vehicle, (B) Gel-50-LN111, and (C) Gel-50-LN121. (Scale bar is 100µm)

While we have demonstrated that laminin containing self-healing hydrogels can be used to enhance cellular retention following suprachoroidal transplantation, we were not able to demonstrate widespread integration of transplanted choroidal endothelial cells. We suspect that this was because the transplant recipients used in this study were disease free (i.e., intact choriocapillaris with no space for new cells to integrate). To further evaluate post-transplant cellular integration, models of choroidal loss with endothelial cell drop out and presence of ghost vessels will be required. While several rodent models of AMD have been reported[70], primary endothelial cell loss is not characteristic in these models. Furthermore, for human cell transplantation, AMD models generated on immunocompromised backgrounds conducive to xenotransplantation will likely be required. To further promote xenograft survival and integration, the hydrogel system developed here can readily be supplemented with additional anti-inflammatory small molecules[71] or vasculogenic growth factors [72], which are designed to suppress the innate immune response and promote endothelial cell repopulation respectively. That said, there is no guarantee that human cells will recognize ques in rodent injury or that the rodent choroidal environment will be conducive to xenograft integration. For these reasons, the immortalized choroidal endothelial cell line used in this study may not be ideal for future development of choroidal cell replacement strategies. Rather, immunologically matched choroidal endothelial cells (generated via iPSC differentiation[64] or direct conversion[73, 74] of host somatic cells) will be used going forward.

In conclusion, in this study we present a library of novel hydrogels generated using laminin, chitosan and dextran via Schiff based chemistry. The developed self-healing hydrogels had desirable mechanical, gelation, and degradation properties, making them ideal for choroidal endothelial cell transplantation. Future studies focused on the use of the developed hydrogels for autologous endothelial cell replacement in animal models of choroidal cell dropout will pave the way for clinical transplantation studies in patients with early stage AMD.

## MATERIALS AND METHODS

### Oxidation of Dextran

Dextran was oxidized using sodium periodate (NaIO_4_) as per previously described protocols [75, 76]. Briefly, 5 grams of Dextran (Sigma-Aldrich, St. Louis, MO) (60 - 76kDa) was dissolved in 200 mL of deionized water. 5 grams of NaIO_4_ (Sigma-Aldrich) was subsequently added and the solution was stirred for 90 mins in the dark. 5mL of ethylene glycol (Fischer Scientific, Waltham, MA) was subsequently added and the solution was stirred for 30 mins to terminate the reaction. The reaction mixture was then dialyzed using MWCO 6-8kDa (Spectra/Por) dialysis tubing for 24 hours in water to remove byproducts. The final product was flash frozen and lyophilized. NMR spectrum was obtained using a Bruker 400Hz NMR system on the oxidized dextran dissolved deuterated water.

### Hydrogel fabrication

Both carboxy methyl chitosan (Biosynth, United Kingdom) and oxidized dextran precursor solutions were prepared using complete endothelial cell growth media. Laminins (BioLamina, Sweden) were used in their undiluted from as provided by the manufacture. 2.5 % w/vol carboxy methyl chitosan and 1% w/vol oxidized dextran were prepared separately and incubated overnight at 37°C. For hydrogel fabrication, laminin and chitosan were mixed together. If choroidal endothelial cells were included, they were resuspended in the previously prepared oxidized dextran solution. Precursor solutions were subsequently mixed and incubated (incubation time was experiment dependent) at 37°C before further use. For suprachoroidal injections, the hydrogel mixture was loaded into a 32G injection device (Hamilton, Reno, NV). For in vitro testing the hydrogel mixture was pipetted into a 6-well plate. For rheological characterization hydrogels were incubated between parallel plates with a solvent trap filled with deionized water.

### Rheological characterization

Rheological measurements are obtained using a 25 mm diameter stainless steel parallel plate geometry, with a solvent trap on the ARES-G2 instrument (TA instruments). To determine gelation time, the hydrogel precursor solutions of laminin, chitosan and dextran were mixed, and 1 mL of the hydrogel solution was immediately pipetted on to the rheometer. A time sweep was performed at 37°C with an oscillatory shear rate of 1 Hz and 1% strain. Prefabricated cylindrical hydrogels, 25 mm in diameter and 2.5 mm in height were used for the storage modulus measurements. The hydrogels were cast in a mold and incubated overnight at 37^0^C. Following incubation hydrogels were carefully placed on the rheometer and low strain frequency sweeps were performed between 1 Hz to 50 Hz at constant axial force of 0.1 N.

### Cell culture

GFP positive choroidal endothelial cells (ICEC2-TS) were cultured in complete endothelial cell growth media (R&D systems) containing 1% primocin (InvivoGen, San Diego, CA). Cells were passaged at a ratio of 1:10 every five days, with complete media changes every other day. During in vitro monitoring of the cell laden hydrogels, the media was changed daily. For in vitro experiments 0.5M cells/ml of the hydrogel were used. For in vivo suprachoroidal injection studies 30k cells/10 ul of the hydrogel or complete media were used.

### Live Dead assay

Calcein AM(4µM) (ThermoScientific, Waltham, MA) was used to label living cells while ethidium homodimer-1 (8µM) was used to label dead cells. Both stains were added to media and incubated for 45 mins before imaging. Live/dead images were obtained using BioTek Cytations 5 (Agilent Technologies, Santa Clara, CA) in aseptic conditions. Media was replaced immediately following imaging to minimize the excess dyes.

### Suprachoroidal injections (animal procedures)

Immune competent Sprague-Dawley rats (1-2 months of age, Charles River, Wilmington, MA) were used to evaluate biocompatibility of the developed hydrogels in vivo. To prevent xenograft rejection immunocompromised SRG rats (1-2 months of age, Charles River, Wilmington, MA) were used as recipients of cell laden hydrogels. Animals were anesthetized using 3-5% inhalant isoflurane gas (Piramal Healthcare, Bethlehem, PA). The eyes were dilated using 1% tropicamide (Alcon Laboratories, Fort Worth, TX). A conjunctival peritomy was created for scleral exposure in the temporal quadrant to help with the suprachoroidal injections as previously described [77–79]. 10 µl of either cell laded hydrogel set in an 32G injection device 0r single cells suspension (unsupported control) were injected via suprachoroidal injections. 2 mg/kg Meloxicam was subcutaneously injected to all the animals following the procedure for pain reduction. Animals were anesthetized using an intraperitoneal dose of 91mg/kg Ketamine and 9.1mg/ kg xylazine. OCT and fluorescent fundus photos were obtained using an in vivo small animal precision ophthalmic imaging instrument (Phoenix Micron IV, Bend, Oregon). Following examination, animals were sacrificed using 150 mg/kg euthasol and the eyes were harvested and fixed in 4% paraformaldehyde.

### Tissue processing

To orient the eyes for embedding, a limbal suture was placed at the site of the injection. The cornea and the lens were removed, and the posterior cup was rinsed in increasing concentrations of sucrose. Eyes were embedded in a 2:1 solution of 20% sucrose to Tissue Tek OCT embedding media, flash frozen, and stored at −80°C. 7 µm thick sections of the posterior cup were taken using a Cryostat Microtome (Leica Biosystems, Wetzlar, Germany)

### Immunostaining

Cryosections were blocked using SuperBlock T20 Blocking buffer (ThermoScientific) for 1 hour. The sections were subsequently incubated in anti-rabbit IBA1 (1:400) overnight at 4°C. Sections were subsequently incubated with Donkey anti Rabit AF488 (1:500) for 1 hour at room temperature. DAPI was used to label cell nuclei and slides were mounted using FluorSave Reagent (Sigma Aldrich) mounting media. Sections were imaged using Biotek Cytations 5 (Agilent Technologies).

