## Supplementary Information for "Development of self-healing hydrogels to support choroidal endothelial cell transplantation for the treatment of early age related macular degeneration"

**Affiliations:** [1]Institute for Vision Research, Carver College of Medicine; University of Iowa, Iowa City, IA; [2] Department of Ophthalmology and Visual Sciences, Carver College of Medicine University of Iowa, Iowa City, IA; [3]Department of Pharmaceutical Sciences and Experimental Therapeutics, University of Iowa, College of Pharmacy, Iowa City, IA; [4]Department of Dermatology, University of Iowa Hospitals and Clinics, Iowa City, IA.

**Keywords:** hydrogels, choroidal endothelial cells, suprachoroidal, cell transplants, self-healing, injectable, xenograft

**Supplementary Table**

| Gel | Carboxy methyl Chitosan (mg/ml) | Laminin (ug/ul) | Dextran (mg/ml) |
|---|---|---|---|
| 1 | 20 | 0 | 2 |
| 3 | 12.5 | 30 | 2 |
| 5 | 7.5 | 50 | 2 |

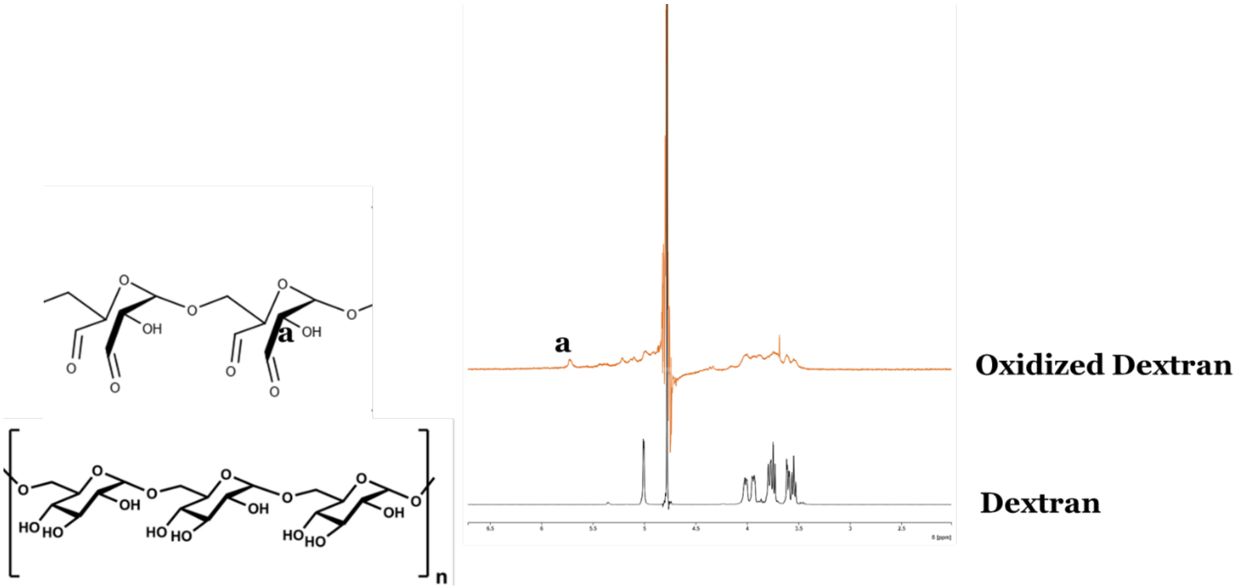


**Figure-S1:** $^1$H NMR spectrum comparing the dextran and oxidized dextran.

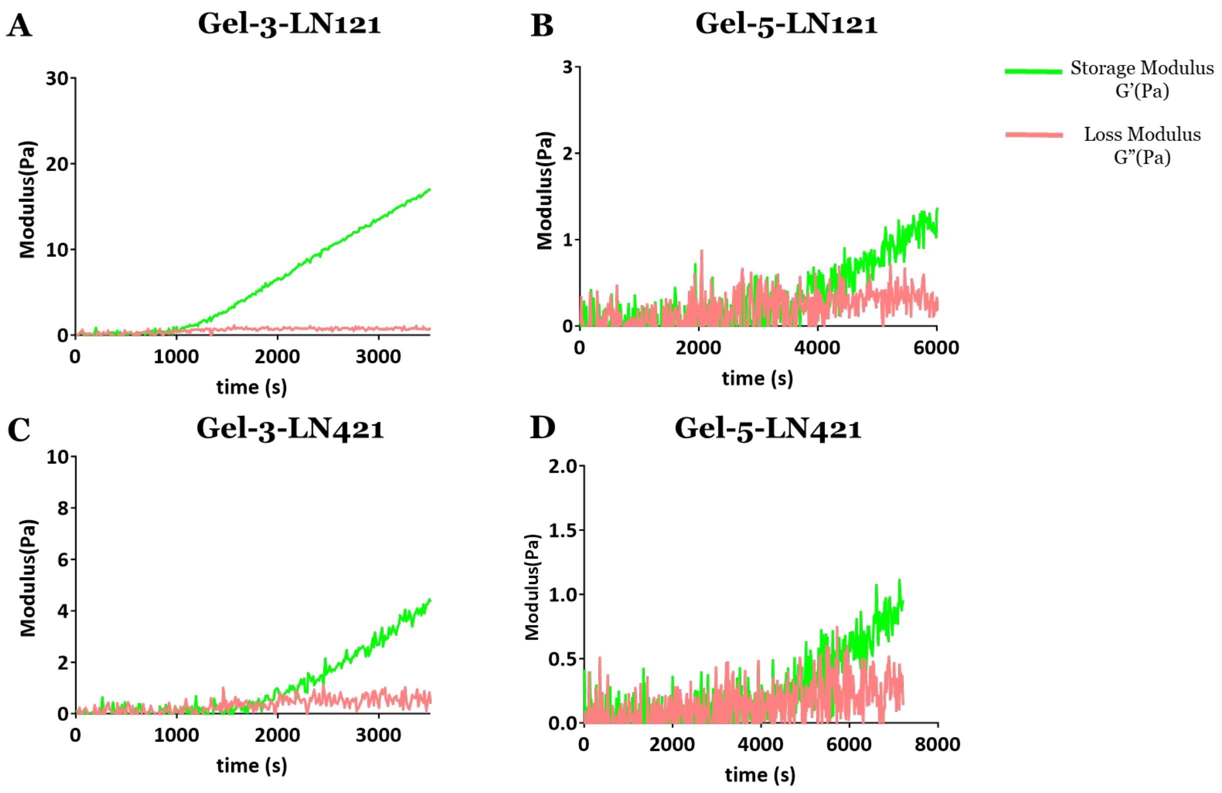


**Figure-s2:** Time sweeps of the A) Gel-3-LN121, B)Gel-5-LN121, C) Gel-3-LN421, and D)Gel-5-LN121 showing the gelation time at the crossover point of the storage(G') and loss modulus(G")

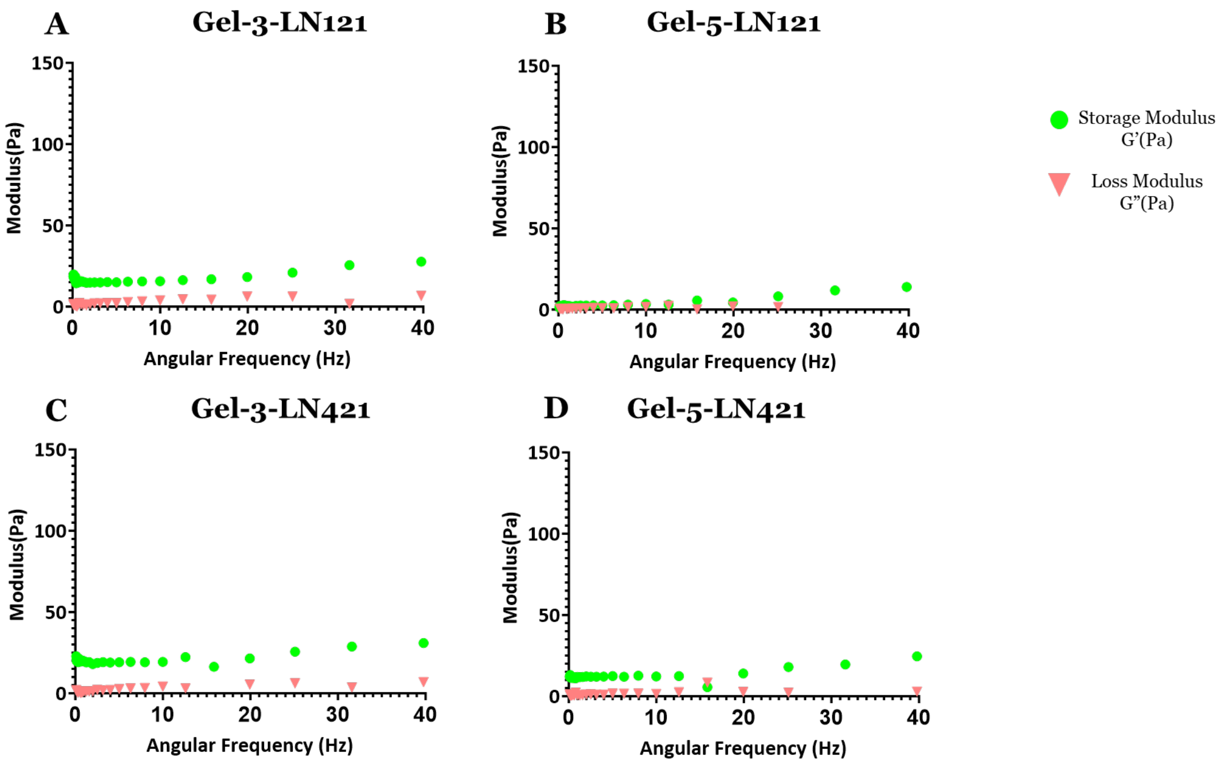


**Figure-S3:** Frequency dependance of the storage modulus (G') and loss modulus (G") of A) Gel-3-LN121, B)Gel-5-LN121, C) Gel-3-LN421, and D)Gel-5-LN121.

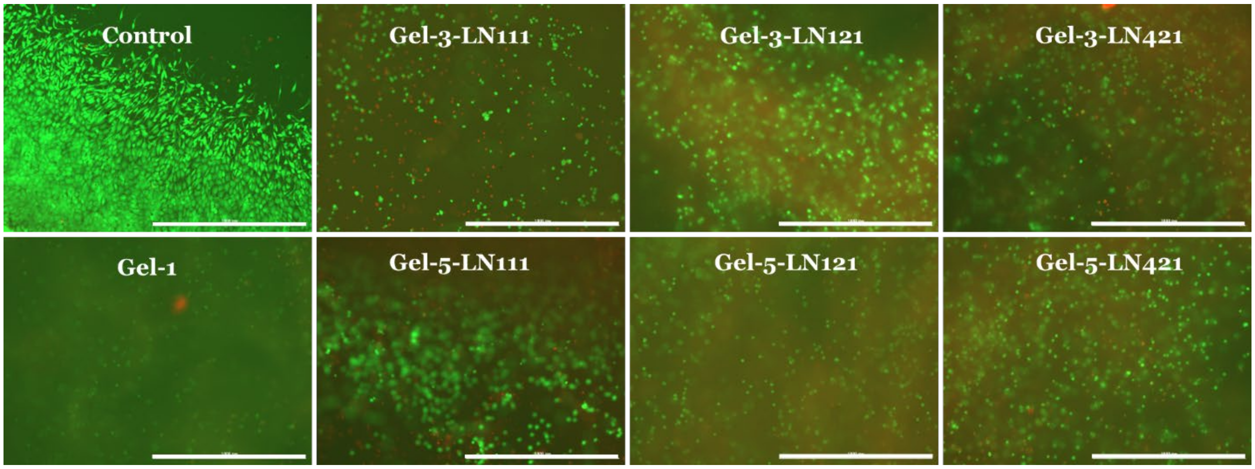


**Figure-S4:** Live dead staining of the cell laden hydrogels synthesized using 3 different isoforms of Laminin (day-7). ICEC2-TS were stained using a live dead stain consisting of calcien-AM as the live stain (green) and ethidium homodimer-1 as the dead stain (red) (Scale bar is 1000μm)

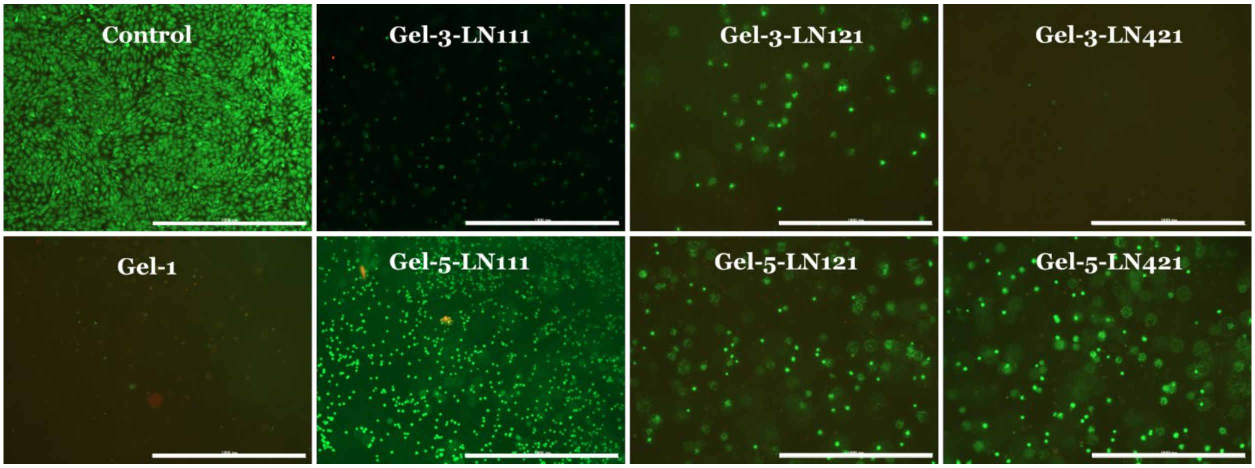


**Figure-S5:** Live dead staining of the different cell laden laminin based hydrogels extruded through 30G fine needle. using 3 different isoforms of Laminin (day-7). ICEC2-TS were stained using a live dead stain consisting of calcien-AM as the live stain (green) and ethidium homodimer-1 as the dead stain (red) (Scale bar is 1000μm)
